# Dual long-axis reorganization of hippocampus in youth

**DOI:** 10.1101/2023.11.03.565423

**Authors:** Debin Zeng, Qiongling Li, Deyu Li, Yirong He, Xiaoxi Dong, Shaoxian Li, Shenghan Bi, Xuhong Liao, Tengda Zhao, Xiaodan Chen, Yunman Xia, Tianyuan Lei, Lianglong Sun, Weiwei Men, Yanpei Wang, Daoyang Wang, Mingming Hu, Zhiying Pan, Shuping Tan, Jia-Hong Gao, Shaozheng Qin, Sha Tao, Qi Dong, Yong He, Xi-Nian Zuo, Shuyu Li

**Author notes:** Corresponding authorship. Address correspondence to Shuyu Li, Xi-Nian Zuo, Yong He.

## Abstract

The reorganization of human hippocampus, especially its interaction with cortex, remains largely undefined in youth. The organization of a single hippocampal long-axis has been predominantly characterized as monotonic^1–6^, despite recent indications of nonmonotonic features in neuron density^7^ and geometric eigenmodes^8^. While the human cortical hierarchy has been well recognized for significant developmental and evolutionary advantages^9–12^, hippocampus has been typically considered an evolutionarily conserved brain structure^1,13,14^, and overlooked regarding its integrative role of cortical hierarchical processing during development. Here, we corroborated the presence and significance of a dual long-axis representation of the hippocampal connectome and geometry including both linear and quadratic gradients along its long-axis in youth. This finding was robust across two independent large-scale developmental cohorts. Charting development of the dual long-axis gradients underscored their specific contributions to the cortical hierarchy maturation from the frontoparietal and salience/ventral attention networks. The observed developmental variability in spontaneous brain activities in youth parallels the gradients of myelin content. During childhood through adolescence to early adulthood, the hippocampus reorganized the dual long-axis by gradually relaxing its geometric constraints on the intrinsic network organization of cortical spontaneous activity for refined executive functions. Molecular processes underlying such reorganization of the dual long-axis in hippocampus are linked to neural growth, stress hormone regulation, and neuroactive signaling. Our findings enrich the understanding of hippocampal-cortical reorganizational principles across structural, functional, and molecular dimensions as well as its maturation, and define the plasticity distribution within the human hippocampus at systems level, holding potentials to enhance and translate neurodevelopment and neuropsychiatric healthcare.

---

The spatial arrangement organization principle of the human brain supports its elaborate functions, a concept akin to that observed in numerous natural systems. For instance, Earth’s climate zones are organized along a gradient of solar radiation reception; elevation gradients in mountains drive changes in species diversity; and animal groups, such as wolves, often exhibit a gradient of social interaction that influences their hierarchical structures. Understanding the organization principles of a system is crucial for comprehending its functioning. The hippocampus, recognized as a pivotal brain structure, follows the same principle and carries notable significance not only concerning its phylogenetic origins^15^ but also its engagement in a broad range of cognitive functions and behaviors beyond memory and spatial navigation^16–23^. In the past decade, a consensus has been growing regarding distinct organizational differentiation along the anterior-posterior axis (i.e., the long-axis) of the hippocampus in various species^1–3,24^. Such differentiation is reflected in its microstructure, gross morphology, connectivity, gene expression, electrophysiological properties, representational granularity, and functional profiles. This anterior-posterior distinction is not characterized by abrupt shifts but rather by a continuous spectrum or gradient that extends along the long-axis, encompassing multiple organizational traits^2,4–6,25,26^. Consequently, investigating the gradient along the long-axis aids in comprehending how the hippocampus processes information and facilitates diverse cognitive functions, along with revealing its implications for various neurological and psychiatric conditions^1,3,24^. Despite significant advancements in elucidating the governing long-axis principle of the hippocampus, most of the literature has proposed a monotonic gradient while a few recent studies have reported organizational characteristics that deviate from this monotonic gradient. Specifically, an investigation leveraged *BigBrain* data to reveal an inverted U-shaped distribution of neuron density along the hippocampal long-axis^7^. Additionally, one primary geometric eigenmode of the adult hippocampus exhibits a quadratic pattern along the long-axis^8^. Hence, the following question has arisen: do the current well-established principle of monotonic long-axis gradient adequately capture the hippocampal-cortical organization?

The unimodal-to-transmodal gradient, known as the hierarchical gradient, stands as a prominent and extensively studied organizational and topographic principle within the cortex^9,10,27–31^. This gradient has been evidenced across various organizational traits, including gene expression, cortical microstructure, connectivity, evolutionary expansion, intrinsic dynamics, development, and functional attributes. This gradient offers a systematic way to understand how these characteristics covary in the cortex and furnish a mechanistic comprehension of the emergence of distributed cortical functional specialization and integration. It can thus serve as an integrated cortical coordinate system relevant to human neuroscience. The dominance of hierarchical processing is higher in humans^9^ and macaques^10^ than in other species such as marmosets^11^ or rodents^32^. The prominence of hierarchical gradients in the human brain was found to be established during adolescence^12,33^. These observations underscore the significant evolutionary and developmental benefits associated with this gradient. However, the hippocampus is widely considered an evolutionarily conserved brain structure, exhibiting similar structural and functional organization across different species, including rodents^1,13,14^. Notably, the hippocampus has also been identified as one of the evolutionary origins of the neocortex^15^. An intriguing inquiry emerges: relying on the hippocampal organization axis, whether, and if so, how this evolutionarily conserved region integrates into cortical hierarchy development.

Recent rodent studies suggest that the hippocampus experiences critical periods of heightened plasticity during childhood and adolescence. These periods make the hippocampus sensitive to environmental experiences, thus sculpting neural networks and establishing the basis for acquiring adult hippocampus-dependent cognitive functions^34–38^. Human studies have uncovered significant distinctions in the developmental trajectory of anterior and posterior hippocampal volume^39,40^, functional activation^41,42^, hippocampal-cortical networks^43–45^, and structural covariance^46^. Furthermore, the development of long-axis subregions differentially relates to hippocampal-relevant learning and memory outcomes, such as memory encoding and retrieval^40,43,45^, as well as associative memory^47^. However, hippocampal organization development is typically studied in discrete, independent subregions along the long-axis, which imposes clear boundaries between parcels and uniformity within them. This approach fails to effectively capture gradual changes and broader spatial relationships, while gradient mapping approaches can be used to identify principal axes of variance in data through embedding techniques, making fewer assumptions about internal organization^4,9^. An analysis of the development of gradient organization in the hippocampus over youth can provide insights into the developmental spatiotemporal brain organization that is crucial for pinpointing positions and periods of heightened plasticity. Developmental profiles of spatiotemporal gradients in the neocortex^28^ and cerebellum^48^ were recently unveiled, but they remain undiscovered for hippocampal development.

Here, we profile the hippocampal-cortical connectivity gradients and their developmental trajectories to demonstrate the dual long-axis gradients in the human hippocampal connectome in youth. Projecting the connectome gradients onto the cortex, we clarified how distinct cortical hierarchies allocate functional connectivity differently along the long-axis, thus coding the hippocampus’s intricate and multifaceted role in cortical hierarchical processing. These discoveries challenge classical views that propose a monotonic gradient of structural and functional differentiation along the hippocampal long-axis, and question the traditional notion of the hippocampus as being evolutionarily conserved in terms of its organization. We observed substantial developmental reorganization of dual long-axis gradients in supporting the maturation of the cortical hierarchy in youth. The reorganization further unfolds that the human hippocampus continues to loosen its geometric gradient constraints on functional gradients to support the executive function performance. Notably, we revealed that neurodevelopmental variability in the functional gradient profiles mirrors a gradient associated with a plasticity-limiting factor (myelin content, estimated by T1w/T2w ratio^49^). At micro-level, we found that neural growth, stress hormone regulation, and neuroactive signaling are involved in this geometry-function-cognition alignment, facilitating such reorganization of the dual hippocampal long-axis gradients in youth.

## Results

Our primary findings were obtained from a large-scale MRI dataset sourced from the Human Connectome Project Development (HCP-D), encompassing 652 typically developing participants aged 5-21 years (351 females). The validation results were obtained from a longitudinal multimodal MRI dataset from the Children School Functions and Brain Development Project (CBD, Beijing Cohort), consisting of 300 healthy children across childhood and early adolescence (140 females, aged 6-13 years, 478 total scans, Supplementary Fig. 1). Structural and resting-state functional MRI (rs-fMRI) underwent HCP minimal preprocessing^50^ with an additional denoising process for fMRI (see the Methods section, Image preprocessing). To capitalize on the advantages of mitigating the partial volume effect and accommodating the diverse variants of hippocampal folding observed among individuals^4,51^, we employed an innovative deep learning technique integrated with a topological constraint approach (HippUnfold^52^) to segment the hippocampus and generate mid-thickness surfaces (483 vertices) featuring coordinates of two intrinsic geodesic axes (posterior-anterior (P-A) and proximal-distal (P-D) axes). This approach effectively captured the unique hippocampal conformation of each subject for subsequent analyses (**Figure 1a**; Supplementary Fig. 2; see the Methods section, Hippocampus segmentation and mid-thickness surface generation).

**Figure 1.**
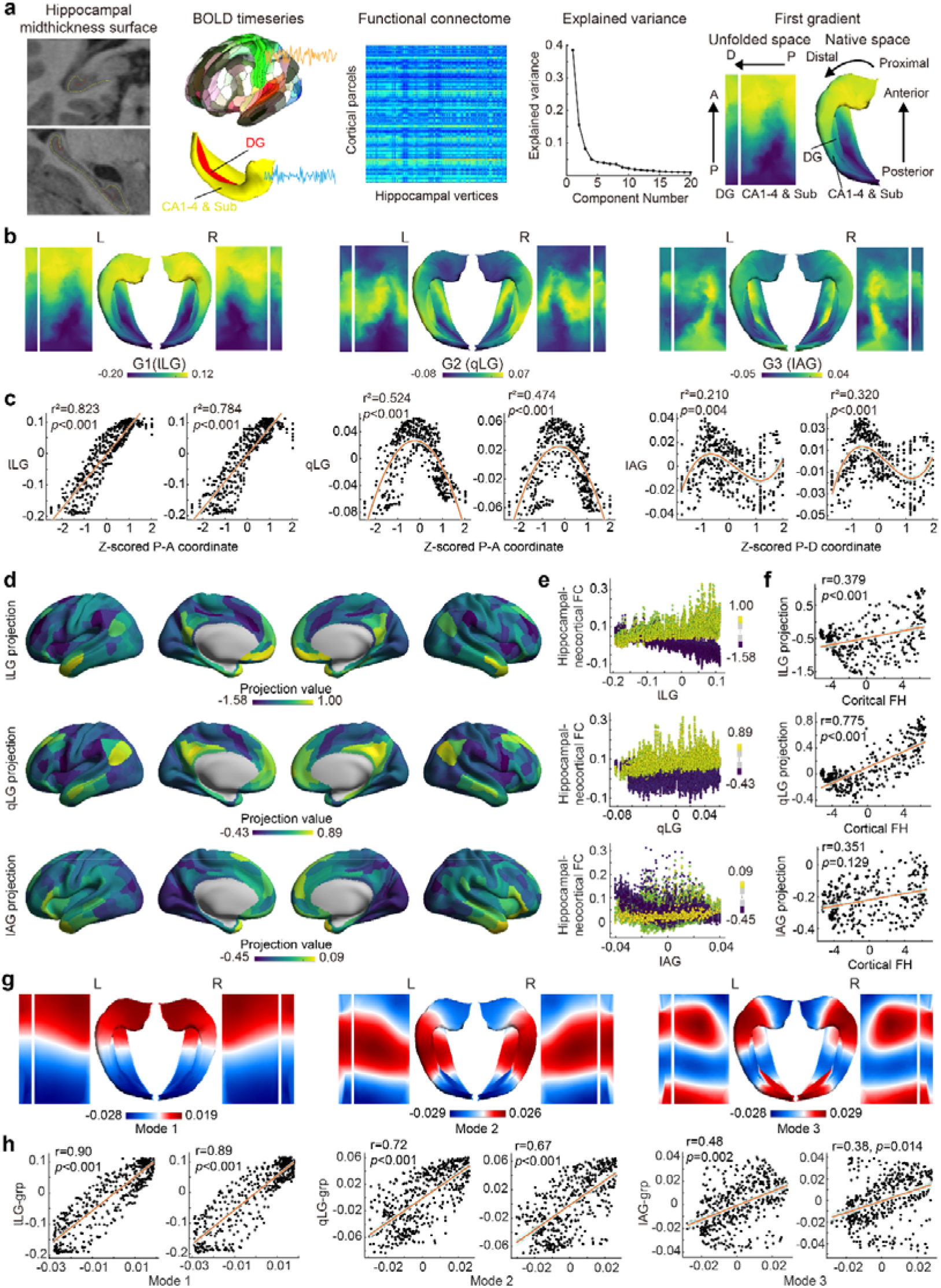
Topographic characterization, cortical projection, and geometric constraints of major hippocampal gradients in youth. **a**, Methodological overview for identifying hippocampal gradients. The gradient’s topography is presented on both the native (folded) and unfolded mid-thickness surfaces, located on the far-right side. The primary geodesic axes, namely posterior-anterior (P-A) and proximal-distal (P-D), are displayed by arrow. On the P-D axis, with the dentate gyrus (DG) representing the most distal region, the coordinates indicate the distance to the neocortex (isocortex), and thus reflect the iso-allocortical axis. Coordinates for each geodesic axis (P-A or P-D) are determined based on the Laplace field across hippocampal gray matter, spanning from one tissue boundary (posterior or proximal end) to the opposite boundary (anterior or distal end). **b**, Topographic characterization of the first three hippocampal gradients at the overall group level. **c**, Relationships between hippocampal gradients and anatomical positions, specifically denoted by P-A and P-D coordinates. **d**, Group-averaged cortical projection of the left hippocampal gradients. **e**, Relationship between left hippocampal gradients and the cortical-hippocampal FCs corresponding to the cortical regions that exhibit the top 5% and bottom 5% projection values. Each data point within the diagram signifies the FC and gradient value corresponding to an individual vertex located on the hippocampal surface. The color indicates the projection values. **f**, Correlation between the projection pattern of specific gradients in the left hippocampus and cortical functional hierarchy (FH). **g**, Group-averaged first three geometric eigenmodes obtained through the Laplace–Beltrami operator. **h**, Relationship between these eigenmodes and hippocampal gradients. Abbreviations: lLG, linear long-axis gradient; qLG, quadratic long-axis gradient; IAG, iso-allocortical gradient.

### Dual long-axis of hippocampal-cortical functional organization in youth

By computing the Pearson correlation between the rs-fMRI time series of each hippocampal vertex and each cortical region (defined by the Glasser atlas^53^), we constructed hippocampal-cortical connectomes that depict the coupling of functional signals between them. We then applied the diffusion embedding method, a nonlinear dimension reduction algorithm, to obtain the hippocampal functional gradient (See the Methods section, Hippocampal feature mapping and gradient computation, **Figure 1a**). The group-level hippocampal gradient was initially derived by averaging the functional connectivity (FC) across all participants (**Figure 1b**) and within several age-specific groups (**Extended Data Fig. 6**) in the HCP-D dataset.

The elbow point in the explained variance plot (**Figure 1a**) indicates that the first five gradients collectively account for a substantial portion of the variance in hippocampal FCs (72.9% and 71.5% for the left and right hemispheres, respectively; refer to **Figure 1b** and **Extended data Fig. 1**). The first principal gradient (G1) explained 39.5% and 38.6% of the variance in the left and right hemispheres, respectively. It exhibited pronounced differentiation along the long axis and demonstrated a strong linear relationship with P-A coordinates (r^2^=0.823, *p*<0.001 for the left hippocampus; r^2^=0.784, *p*<0.001 for the right hippocampus; see the left panel of **Figure 1b and c**). Conversely, there was no significant evidence of an association between this gradient and the P-D coordinates that indicated the distance to the neocortex (**Extended Data Fig. 2**). Consequently, we have designated this gradient as the linear long-axis gradient (lLG). Innovatively, the second-order gradient (G2), explaining 16.6% and 15.5% of the variance in the left and right hemispheres, respectively, exhibited a distinctive pattern of transition from the hippocampal body to the head and tail (see the middle panel of **Figure 1b**). The G2 showed a significant quadratic relationship with the P-A coordinate (r^2^=0.524, *p*<0.001 for the left hippocampus; r^2^=0.474, *p*<0.001 for the right hippocampus; middle panel of **Figure 1c**). Conversely, we found no convincing evidence of any association between this gradient and P-D coordinates (**Extended Data Fig. 2**). Therefore, we have termed this gradient the quadratic long-axis gradient (qLG). Furthermore, the third-order gradient (G3), explaining 7.7% and 8.3% of the variance in the left and right hemispheres, respectively, exhibited a P-D differential pattern (see the right panel of **Figure 1b**). This relationship was supported by a significant cubic association between the G3 and the P-D coordinates (r^2^=0.210, *p*=0.004 for the left hippocampus; r^2^=0.320, *p*<0.001 for the right hippocampus; see the right panel of **Figure 1c)**, while little credible evidence shows the relationship between the G3 and the P-A coordinates (**Extended Data Fig. 2**). This gradient exhibits similarities to the previously established cytoarchitectonic variations within the hippocampus^1,3^. Hence, we named this gradient the iso-allocortical gradient (IAG). We refrained from analyzing G4 and G5 due to the lack of clarity regarding their biological significance. The statistical significance of the correlations between gradients and anatomical positions was evaluated using a spatial null permutation test^54^ (see the Methods section, Statistical testing of spatial alignment). Notably, these topographic profiles of the first three gradients and their relationships with the anatomical geodesic axis were also consistent on the independent validation dataset (CBD), and the corresponding results can be found in **Extended Data Fig. 3**.

We aimed to further investigate how the multifaceted organization of the hippocampus supports large-scale integration into cortical networks. We projected the principal hippocampal gradients onto the cortex through the hippocampal FC corresponding to each cortical region (see the Methods section, Cortical projection of hippocampal gradients). We performed this analysis for the first three gradients for each participant and computed the group-averaged projection map (see **Figure 1d** for the group-averaged projection of the left hippocampal gradient and **Extended Data Fig. 4** for the right). **Figure 1e** illustrates the relationship between the hippocampal gradient and the cortical-hippocampal FCs corresponding to the cortical regions that exhibit the top 5% and bottom 5% projection values (left hippocampus; see **Extended Data Fig. 4** for the right). As anticipated, the acquired projection values effectively distinguished the cortical-hippocampal FC patterns across distinct cortical regions, with varying projection values for a given hippocampal gradient displaying differing allocations of FCs along that hippocampal gradient. Specifically, the top panel in **Figure 1e** shows that higher lLG projection values signify a heightened level of connectivity with the positive lLG region (anterior hippocampus) compared to the negative lLG region (posterior hippocampus). Similarly, the middle panel in **Figure 1e** illustrates that higher qLG projection values correspond to stronger connectivity with the positive qLG region (hippocampal body), in contrast to the negative qLG region (hippocampal head and tail). Significantly, we established a strong correlation between the projection pattern of specific gradients and a well-established cortical functional hierarchy (FH) map^9^. The projection distributions of the lLG and qLG, with a particular emphasis on the qLG, exhibited significant correlations with the cortical FH (lLG projection: left hippocampus r=0.379, *p*<0.001, right hippocampus r=0.381, *p*<0.001; qLG projection: left hippocampus r=0.775, *p*<0.001, right hippocampus r=0.788, *p*<0.001; IAG projection: left hippocampus r=0.351, *p*=0.129, right hippocampus r=0.426, *p*=0.082). Refer to **Figure 1f** (left hippocampus) and **Extended Data Fig. 4** (right hippocampus) for detailed visualizations. Statistical significance was assessed using the same spatial null permutation test mentioned before. Importantly, the observed associations between the dual long-axis gradient projection and cortical FH were consistently found in the independent CBD dataset, as depicted in **Extended Data Fig. 5**. Our study demonstrated that the hippocampus actively integrates into cortical hierarchical processing. Notably, our observations indicated that the qLG plays a particularly crucial role in the cortical hierarchy, surpassing the importance of other gradients. This finding is particularly intriguing considering the widely held belief that the hippocampus is evolutionarily conserved, while cortical hierarchical processing is observed to be more dominant in humans and macaques^10^ than in other species, such as marmosets^11^ or rodents^32^. These observations suggested that human hippocampal dual long-axis gradients, closely intertwined with cortical hierarchical processing, possess noteworthy evolutionary advantages, thereby prompting a reevaluation of classical perspectives.

Geometric constraints characterized by Laplace eigenmodes were recently considered dominant factors in shaping the functional organization of neocortical and non-neocortical structures, offering a simpler and mechanistically informative framework^8^. We derived the geometric eigenmodes of the hippocampus for each participant using the Laplace– Beltrami operator (we discarded the first constant mode, see Derivation of hippocampal geometric eigenmodes in the Methods section). **Figure 1g** showcases the group-averaged first three eigenmodes. Our analysis revealed a significant correlation between these geometric modes and the first three functional gradients, thereby establishing a strong association between hippocampal geometry and dynamics in youth. This relationship is demonstrated in **Figure 1h**, where the lLG exhibited a correlation with Mode 1 (left hippocampus: r=0.90, *p*<0.001; right hippocampus: r=0.89, *p*<0.001), the qLG with Mode 2 (left: r=0.72, *p*<0.001; right: r=0.67, *p*<0.01), and the IAG with Mode 3 (left: r=0.48, *p*=0.002; right: r=0.38, *p*=0.014). These results underscored the fundamental role of geometry in shaping hippocampal spontaneous functional activity in youth.

Collectively, these findings provided compelling evidence for the presence of dual long-axis differentiation in the human hippocampus across the functional connectome, in their geometry, and in their roles in cortical hierarchical processing. We revealed that distinct cortical hierarchies allocate FC differently along hippocampal gradients, particularly for the qLG. These discoveries challenge conventional views that posit a monotonic gradient of structural and functional differentiation along the long-axis of the hippocampus, and call into question the wide belief of the hippocampus as an evolutionarily conserved structure according to its structural and functional organization.

### Dual long-axis gradients develop parallel to myelin maturation

To visually examine the age-related alterations in hippocampal gradients across childhood, adolescence, and young adulthood, we computed the gradients of averaged connectivity within four age-specific groups (5-9, 10-13, 14-17, 18-21 years) comprising 111, 201, 188, and 152 participants, respectively. Our findings indicated that the distribution of the lLG exhibited expansion during childhood, followed by a subsequent contraction upon entering adolescence. Conversely, the distribution of the qLG continued to expand throughout this developmental period while the IAG displayed relative stability within this timeframe (**Figure 2a**, **Extended Data Fig. 6 and 7a**).

**Figure 2.**
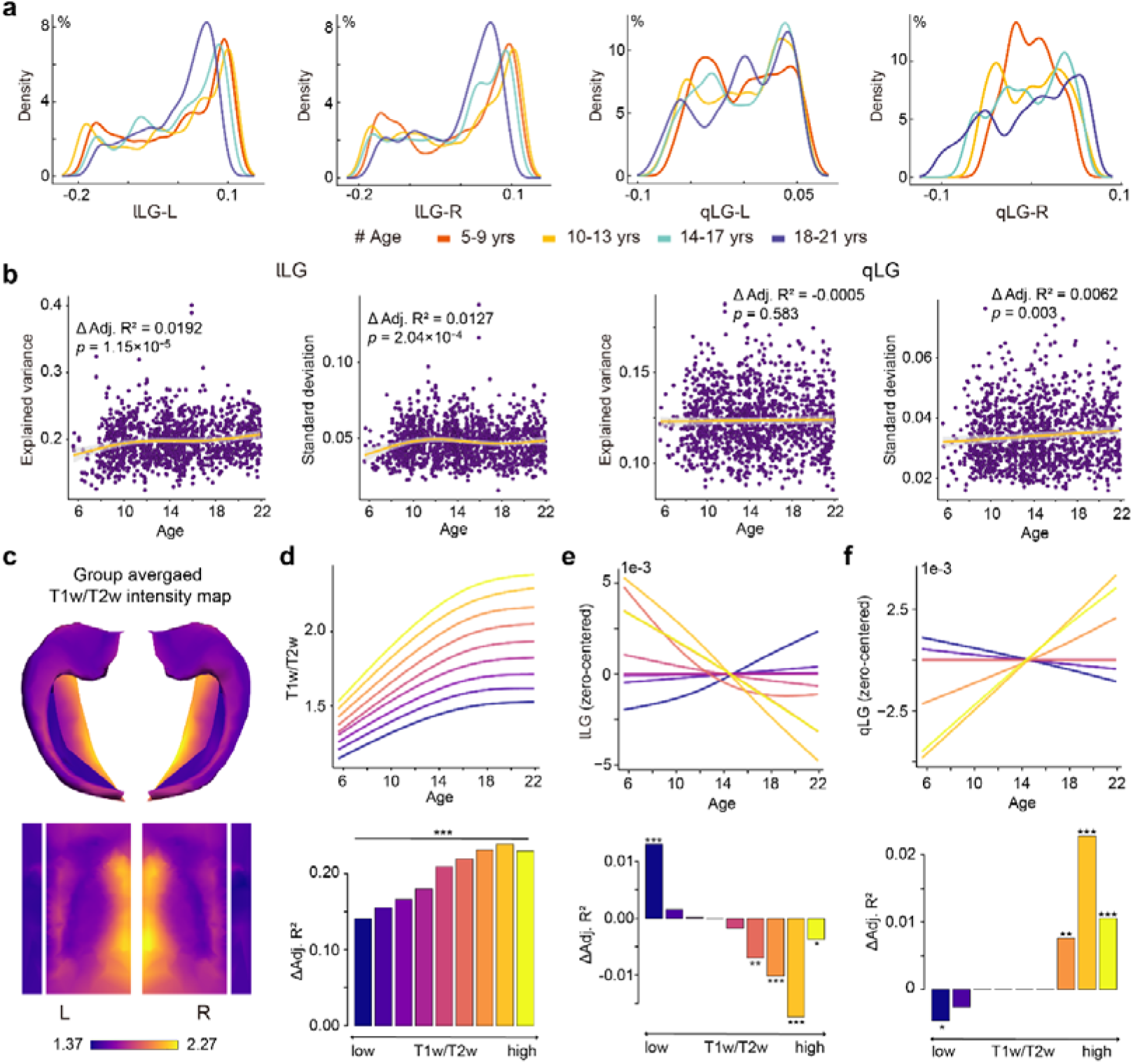
Development of dual long-axis dimensions of hippocampal gradient parallels myelin maturation. **a**, Density estimation of dual long-axis gradient values within four age-specific groups (5-9, 10-13, 14-17, 18-21 years). **b**, The developmental progression of the explained variance and standard deviation of dual long-axis gradients. **c**, Group-averaged T1w/T2w intensity map. **d**, Developmental trajectories and age effects (Δ Adjusted R^2^) of the average T1w/T2w intensity within each of the nine uniform bins along the group-averaged T1w/T2w intensity axis. Similarly, **e** and **f** depict the corresponding developmental patterns for lLG and qLG within the same nine bins, respectively. All the developmental trajectories in **e** or **f** are zero-centered to facilitate comparison between them. Note: * *p*<0.05, ** *p*<0.01, *** *p*<0.001, FDR corrected.

In addition, we examined the developmental trajectories of explained variance and distribution characteristics, including standard deviation, range, skewness, and kurtosis, corresponding to these gradients across all participants using generalized additive models (GAMs) (**Figure 2b**, **Extended Data Fig. 7b and c**; the displayed confidence band is associated with a 95% confidence level). We fit the model with a smooth term for age as a fixed effect; sex, scanning site, and in-scanner head motion as linear covariates; and hemisphere as a random effect. We calculated the variance explained by age (Δ Adjusted R^2^) as the effect magnitude and determined the direction of the age effect by the sign of the average derivative of the age fit (see the Methods section, Analysis of developmental effects). Our results revealed a progressive dominance of the lLG, as indicated by an increase in explained variance (Δ Adjusted R^2^ = 0.0192, *p* = 1.15 X 10^−5^). Furthermore, the trajectory of standard deviation for lLG indicated significant changes, with an increase during childhood (5-11 years), a decrease during adolescence (12-17 years), and subsequent stability in adulthood (18-21 years) (Δ Adjusted R^2^ = 0.0127, *p* = 2.04X 10^−4^). The qLG demonstrated increasing differentiation throughout the entire period, as evidenced by the developmental trajectory of the standard deviation (Δ Adjusted R^2^ = 0.0062, *p*=0.003). The variance explained by the qLG remained relatively stable during this period (Δ Adjusted R^2^ = −0.0005, *p*=0.583). Last, we observed reduced dominance of the IAG (Δ Adjusted R^2^ =-0.0047, *p*=0.011). The other distribution characteristics for the first three functional gradients are presented in **Extended Data Fig. 7b and c**.

Considering the pivotal role of myelination in regulating and restraining neural plasticity^55,56^, our investigation sought to elucidate the potential relationship between the maturation of myelin content and the development of hippocampal gradient profiles. Initially, we partitioned the group-averaged T1w/T2w intensity (a structural MRI measure sensitive to myelin content; **Figure 2c**) axis into nine uniform bins to ensure consistency. Then, within each bin, we examined the developmental trajectories of mean myelin content and mean gradient values for each of the first three gradients for all participants. The magnitude and direction of age effects were derived using the same method mentioned before. Consistent with our expectations, myelin content increased continuously throughout childhood and adolescence, reaching a relatively stable level in all bins when individuals entered adulthood. Furthermore, the age effect demonstrated a gradient along the myelin axis, indicating that hippocampal regions with higher myelin content exhibited more pronounced age-related changes in myelin content (**Figure 2d**, **Extended Data Table 1** provides the Δ Adjusted R^2^ and FDR-corrected *p* value for each bin). Interestingly, the developmental variability of the hippocampal dual long-axis gradients also displayed a prominent pattern aligned with the myelin axis (see **Figure 2e**, **f** as well as Extended Data Table 1). Through comparison of the age-related changes in functional gradients with those changes in myelin content across all bins, we identified a significant spatial and temporal correspondence between the refinement of myelin content and the maturation of the functional gradient. Notably, these findings remained consistent regardless of the number of bins used (see **Extended Data Fig. 8 and Supplementary Tables 1-4**).

Collectively, these results showed the significant development of dual long-axis differentiation in the human hippocampus in youth, and suggest a potential association between the development of the hippocampal-cortical FC profile and the maturation of a key regulator (i.e., myelination) involved in developmental plasticity.

### Dual long-axis gradients differently contribute to cortical maturation

Given the observation that dual hippocampal long-axis gradients contribute to the cortical hierarchy, we aimed to further investigate whether the development of lLG and qLG projections is linked to the maturation of the cortical hierarchy. To achieve this, we calculated the correlations between individual gradient projections and the previously mentioned adult cortical FH map. Subsequently, we examined the developmental pattern of this association using the aforementioned GAMs. Remarkably, we observed a strengthening link between both lLG and qLG projections and the cortical FH map as individuals aged, with a particular emphasis on the qLG (lLG: Δ Adjusted R^2^ = 0.008, *p*=0.003; qLG: Δ Adjusted R^2^ = 0.020, *p*=3.75 X 10^−7^; see **Figure 3a and e**). This indicates that the development of differentiation in hippocampal-cortical connectivity along the lLG and qLG axes, especially for the qLG, contributes to the maturation of the cortical hierarchy.

**Figure 3.**
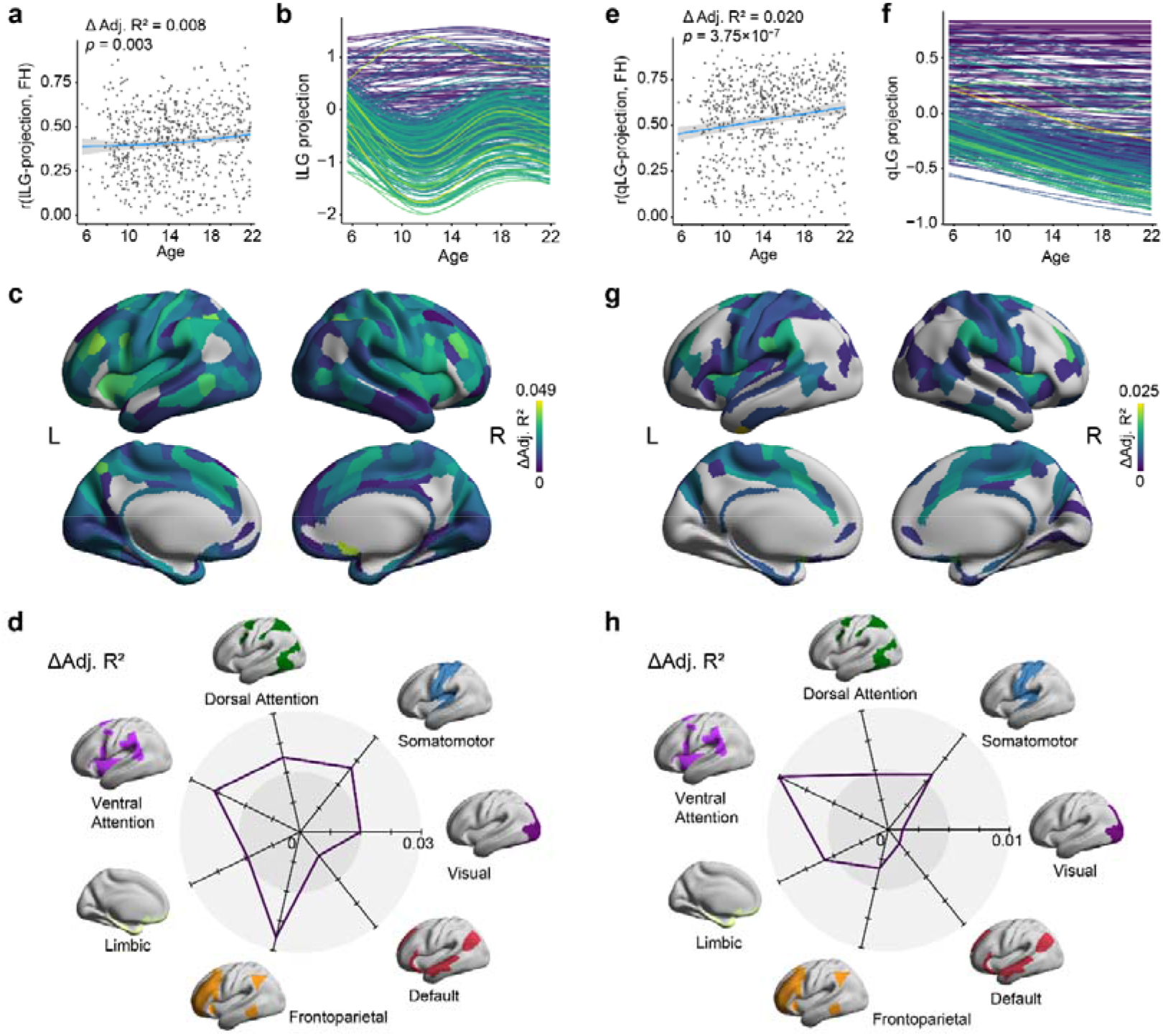
Cortical projection of the hippocampal gradient links to the maturation of the cortical hierarchy in youth. **a**, Development of the association between the hippocampal lLG projection and the cortical functional hierarchy (FH) map. **b**, Region-specific trajectories of hippocampal lLG projections, with each trajectory corresponding to specific cortical regions as delineated by the Glasser atlas. The color of the developmental trajectories was assigned based on the unsigned age effect, which is the same in panel **c**. **c**, Cortical distribution of significant developmental effects (FDR-corrected *p*<0.05) for hippocampal lLG projections. **d**, Distribution of developmental effects within seven intrinsic functional systems defined by Yeo *et al*.^57^. **e**, **f**, **g**, and **h** are similar to **a**, **b**, **c**, and **d** but correspond to the qLG projection.

We also characterized the maturational changes in the hippocampal gradient projections in individual cortical regions. The developmental trajectories of the lLG projection and corresponding significant age effects (FDR-corrected *p*<0.05) are depicted in **Figure 3b and c**. The age effects were unsigned due to the intricate nonlinear nature of the developmental trajectories. Consistent with the development of the lLG range shown in **Extended Fig. 7b**, the range of the lLG projection exhibited an increase during childhood (5-11 years), a decline during adolescence (12-17 years), and subsequent stability in adulthood (18-21 years). We observed that the cortical regions demonstrated significant developmental changes in their cortical-hippocampal FC patterns along the lLG axis. These changes were particularly prominent in regions associated with the frontoparietal and ventral attention systems (**Figure 3c and d**).

Moreover, our investigation revealed that the cortical regions with larger developmental effects (i.e., larger Δ Adjusted R^2^) showed an approximately cubic association with age (**Figure 3b**, green and yellow trajectories), indicating the presence of two inflection points. Intriguingly, one inflection point was evident around the onset of adolescence (approximately 12 years), while the other was observed around the onset of adulthood (approximately 18 years). During childhood, most of these cortical regions increasingly exhibited a preference for connectivity with the posterior hippocampus, which is indicated by the decrease in projection values; however, this trend reversed upon entering adolescence, only to reverse once again upon entering adulthood. This phenomenon potentially regulates the engagement of distinct subregions along the long-axis throughout various developmental stages, tailored to relevant task demands. This hypothesis is in line with several earlier studies revealing age-related disparities in activation along the long-axis during memory encoding and retrieval tasks in youth^42,58,59^. Our findings indicated that there is a significant reorganization of the large-scale integration of the hippocampal long-axis during youth development. The presence of inflection points around the onset of adolescence and adulthood suggests important developmental milestones in the maturation of the hippocampus and its integration with the cortex.

Furthermore, we conducted an additional investigation into the development of qLG projections. As anticipated, the qLG projection range exhibited a consistent and gradual increase throughout the entire period (**Figure 3f**), in line with the findings from the qLG range shown in **Extended Fig. 7b**. Similarly, we observed significant developmental changes predominantly within the ventral attention system, with the age effect being stronger than that of other systems (**Figure 3g and h**). These results from the lLG and qLG indicated that the reorganization of connectivity between both the frontoparietal and ventral attention systems and the dual hippocampal long-axis plays a critical role in the contribution of the hippocampus to the maturation of the cortical functional hierarchy. Specifically, the ventral attention system is of particular importance, as the qLG indicates its greater significance in the cortical hierarchy compared to other gradients. These findings supported prior evidence that the developmental shift in the cortical connectivity gradient, from visual-sensorimotor to hierarchical alignment, is largely driven by connectivity changes in the ventral attention system^60^. Another two studies highlighted the vital role of attention and frontoparietal control systems in cortical hierarchical information flow, reinforcing our findings^61,62^.

Taken together, these results indicate that the development of dual hippocampal long-axis gradients plays an important role in the maturation of the cortical hierarchy, which is largely driven by the development of hippocampal connectivity with the ventral attention and frontoparietal systems. In particular, the FC pattern observed in these systems along the hippocampal lLG axis indicated significant developmental milestones in the maturation of the hippocampus and its integration with the cortex during the transition from childhood to adolescence and from adolescence to adulthood.

### Dual long-axis gradients relax geometric constrains to support the maturation of executive function

We aimed to further investigate the development of geometric constraints on the dual long-axis functional gradients of the hippocampus in youth building on the above findings of significant geometry-function associations. We investigated the development of the coupling between hippocampal geometric modes and their closely correlated dual long-axis gradients. We employed GAMs to fit the age-related trajectory of the coupling, measured using Pearson’s correlation, between these geometric modes and their corresponding gradients (**Figure 4a and c**). Interestingly, the structure-function coupling of the lLG demonstrated a continuous decrease during this period (Δ Adjusted R^2^=-0.055, *p*<1X 10^−16^), whereas the coupling of the qLG displayed a decline with a slight fluctuation (Δ Adjusted R^2^=-0.007, *p*=0.026).

**Figure 4.**
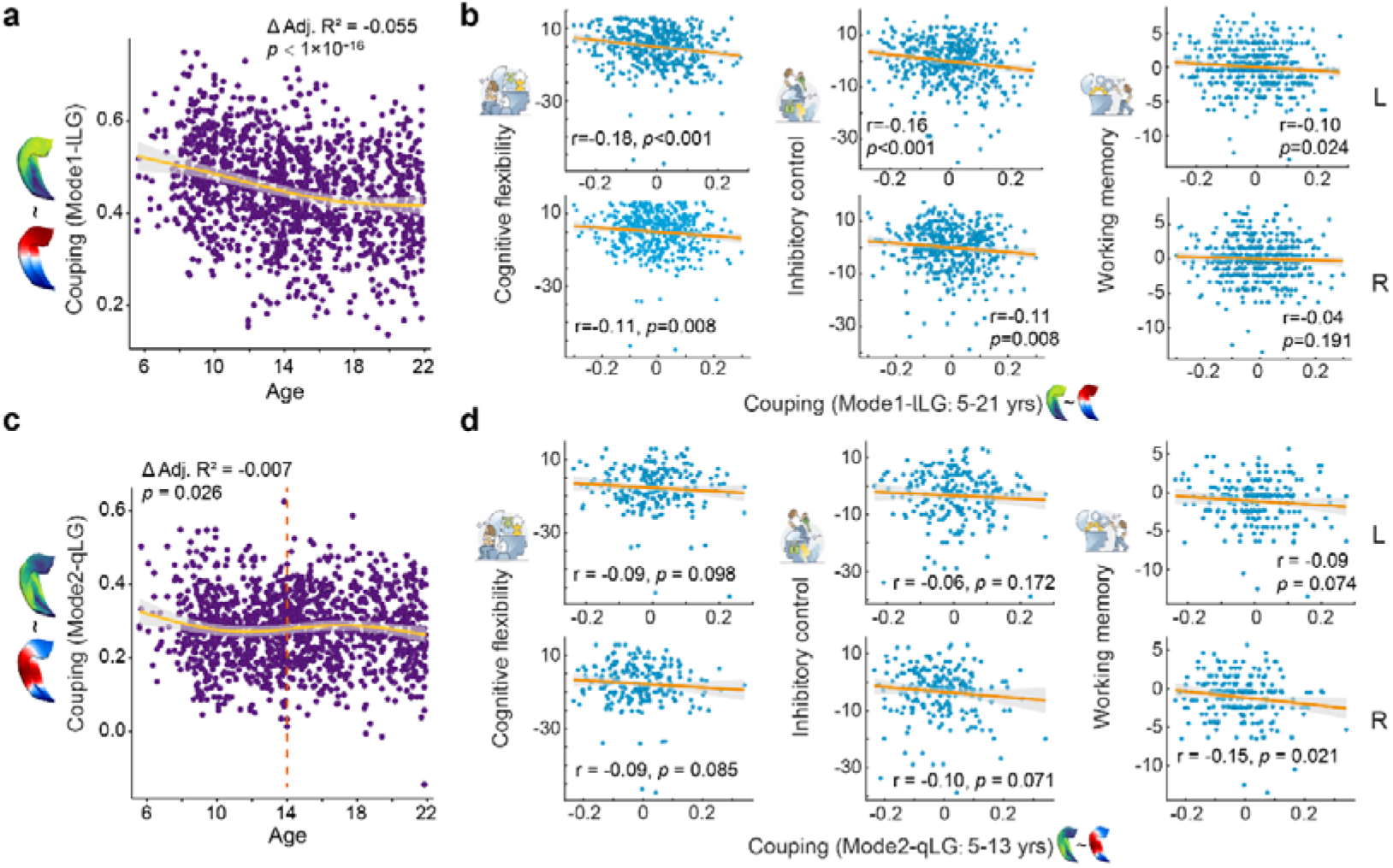
Hippocampal geometric constrains on functional gradients and their association with executive function during youth. **a**, Developmental trajectories of structure-function coupling (Pearson correlation) between geometric eigenmode 1 and the corresponding lLG. **b**, Relationship between the structure-function coupling of the lLG and executive function (EF) performance across the whole developmental period, encompassing cognitive flexibility, inhibitory control, and working memory. Sex, scanning site, and head motion effects were controlled in this analysis. **C,** similar to **a** but corresponds to the structure-function coupling of the qLG. **d**, Relationship between structure-function coupling of the qLG and performance in EF across individuals before 14 years old.

These findings indicated that the spontaneous functional activity of the hippocampus and its integration into the large-scale cortex are significantly dominated by the geometric shape of the hippocampus in youth. Moreover, the observed decrease in structure-function coupling during this specific period suggests the maturation of long-range connectivity within the hippocampus or the more intricate cortical integration with the dual hippocampal long-axis. This maturation process facilitates efficient communication both within and outside the hippocampus and supports more complex hierarchical cortical processing, which ultimately may support the maturation of high-order cognitive functions, such as executive function (EF), a family of top-down mental processes that may be supported by cortical hierarchical organization^63^. EF undergoes a protracted developmental trajectory that parallels the period under investigation. The core and basic EFs include cognitive flexibility, inhibitory control, and working memory. These competencies are integral for promoting mental and physical well-being, as well as fostering cognitive, social, and psychological development^64,65^.

To investigate whether the decrease in structure-function coupling contributes to the performance of core EFs (see the Methods section, Executive function test), we analyzed the association between them. We controlled for sex, scanning site, and head motion effects in this analysis. Importantly, we demonstrated a significant relationship between structure-function coupling of the lLG and core EFs performance across all included ages in the following (**Figure 4b**): for cognitive flexibility, the correlation coefficients were −0.18 (*p* < 0.001) for the left hippocampus and −0.11 (*p* = 0.008) for the right; for inhibitory control, the correlation coefficients were - 0.16 (*p* < 0.001) for the left hippocampus and −0.11 (*p* = 0.008) for the right; regarding working memory, the correlation coefficients were −0.10 (*p* = 0.024) for the left hippocampus and −0.04 (*p* = 0.191) for the right. Interestingly, we also found a significant association between the structure-function coupling of the qLG in the right hippocampus and working memory performance across 5-to 13-year-old individuals (r=-0.15, *p*=0.021), which is distinct from that of the lLG. We conducted this analysis within different age groups because of the non-monotonic developmental trajectory in structure-function coupling of the qLG. All significance levels were assessed using permutation tests. These results suggested a potential link between the decoupling of function from geometry, and the performance of core EFs, and exhibited varying associations across dual long-axis gradients. These results further supported the inference that such decoupling may enhance efficient communication and facilitate more complex hierarchical cortical processing.

### Neuroactive signaling, stress hormone regulation, and neural growth potentially facilitate the dual long-axis reorganization in youth

We conducted a transcriptomic association analysis and developmental enrichment analyses to investigate the neurobiological basis of the dual long-axis functional gradient development. Our primary objective was to explore the presence of gene expression variations along the dual long-axis gradients and identify the key genes involved in this molecular organization. We obtained normalized gene expression data from 58,692 probes, extracted from 173 hippocampal samples of six deceased human donors provided by the Allen Human Brain Atlas. We then employed a LASSO-PCR algorithm to predict each sample’s gradient value using its gene expression profile (see the Methods section, Transcriptomic association analysis of the hippocampal gradients). Through 10 repeated ten-fold cross-validation (CV), the model elucidated an average of 45% (42-48%) variance explication within the lLG (see **Figure 5a and b**) and an average of 7.1% (3.5-10.0%) variance explication within the qLG (see **Extended Data Fig.9a** and **b**). Considering the possible uncertainties tied to model weights for individual probes, which could impact their interpretation, we removed the top 50 positively associated and bottom 50 negatively associated probes iteratively until all 58,692 were eliminated, and reevaluated the model’s CV accuracy after each removal (**Figure 5c**). Notably, removing the initial set of 200 probes (Set 1) resulted in a distinct and unrecoverable decline in CV accuracy, underscoring the significance of this gene set for the model’s performance. Following the removal of the succeeding 500 probes (Set 2; ranked 201–700), a gradual and fluctuating decrease in CV accuracy was observed, and eventually bottomed out (**Figure 5c**, **d**). Conversely, the stepwise removal of sets of 100 random probes resulted in a gradual and intermittent decline in accuracy, culminating at its lowest point solely upon the exclusion of a substantial majority of probes (**Figure 5c**). Additionally, we performed a similar feature deconstructing process for the qLG prediction (**Extended Data Fig.9c**). The results indicated that the removal of important genes did not lead to an irrecoverable decline and showed no significant difference compared to the random gene removal process. Due to the relatively low predictive accuracy and the inability to obtain a robust set of genes for the qLG, our focus centered around the molecular foundation of the lLG.

**Figure 5.**
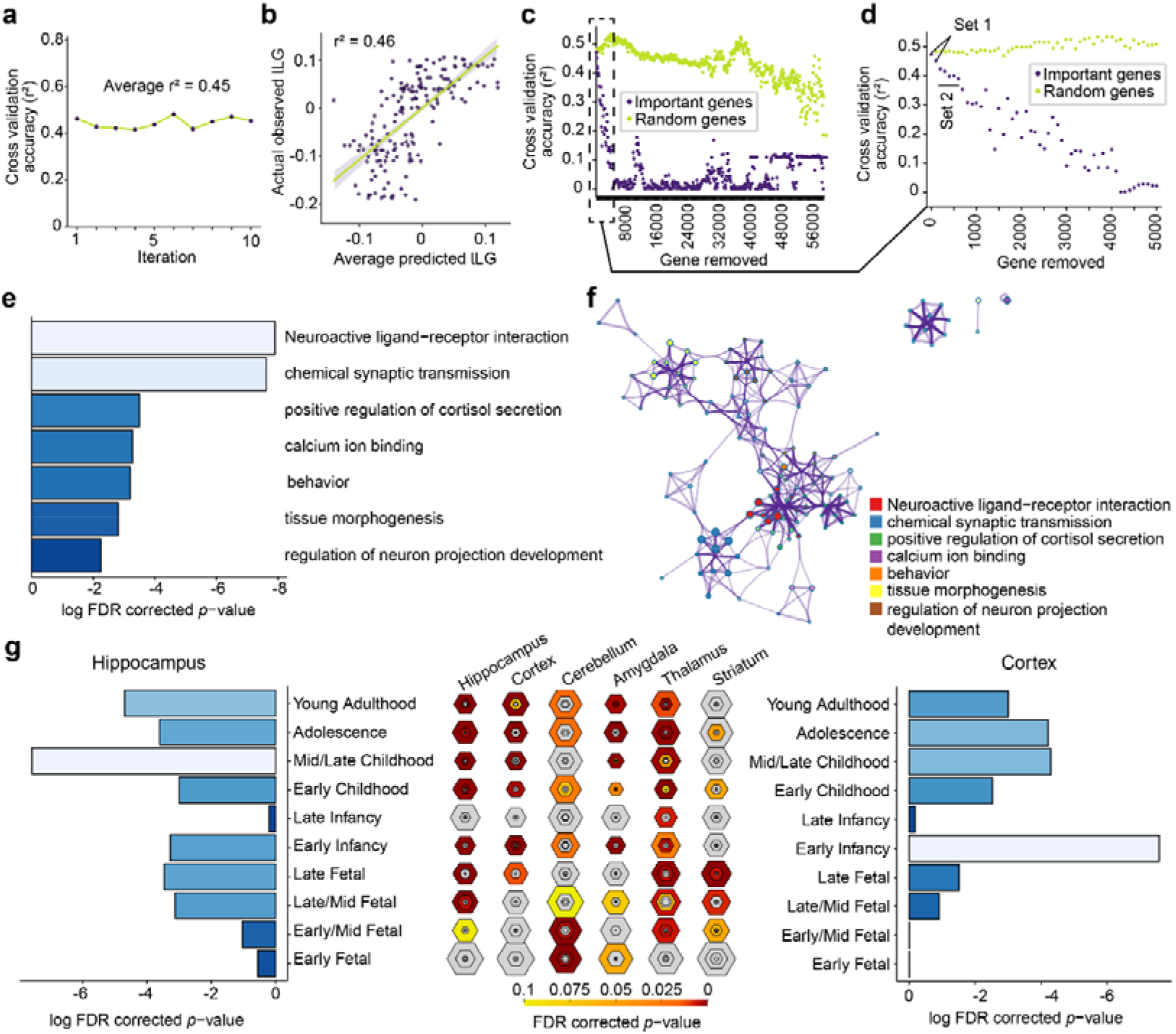
Transcriptomic association analysis of the hippocampal lLG. **a,** Accuracy (r^2^) for ten separate 10-fold cross-validated LASSO-PCR models for lLG prediction using normalized gene expression data provided by the Allen Human Brain Atlas. **b**, Relationship between the average predicted lLG for the aforementioned ten separate 10-fold cross-validated models and the actual observed lLG. **c**, Systematic removal of the top 100 critical probes identified by our prediction model was undertaken. Following each removal, the 10-fold cross-validation accuracy in predicting lLG values was documented (purple dots). To establish a comparative baseline, an iterative process involving the removal of 100 random probes was also executed (chartreuse). **d**, In the zoomed-in view of panel **c**, we scrutinized the initial 50 rounds of 100-probe removal. Notably, a significant decline in performance became evident and this decline was not recovered following the removal of 200 and 700 genes. **e**, A bar graph depicts the enriched biological pathways and gene ontology terms linked to gene Set 1, featuring FDR-corrected *p* values below 0.01. The color scheme represents FDR-corrected *p* values, with lighter shades indicating lower p values. **f**, An enrichment network was constructed by depicting each enriched term as a node and establishing connections between pairs of nodes exhibiting Kappa similarities^66^ surpassing 0.3. The nodes were assigned colors to signify their respective cluster affiliations. **g**, Developmental enrichment analysis. Within the middle panel, the hexagon ring size signifies the proportion of genes within Set 1 that are specifically expressed in a particular brain structure during a specific developmental stage. Hexagon sizes range from least specific (outer hexagons) to most specific (center hexagons) based on varying specificity index threshold (pSI) stringencies (pSI = 0.05, 0.01, 0.001, and 0.0001)^67^. The left and right panels showcase bar graphs illustrating the log-transformed FDR-corrected *p* values for the hippocampus and cortex, respectively.

To delve into the biological processes and molecular functions associated with the pivotal gene set responsible for predicting the lLG, we performed enrichment analyses for biological pathways and gene ontology utilizing Metascape^68^. **Figure 5e** shows the most significant enrichment term clusters derived from the Set 1 list, with FDR-corrected *p* values below 0.01. We adopted enrichment networks by connecting enriched terms with Kappa similarities^66^ exceeding 0.3 to depict similarities between term clusters and redundancies within clusters (**Figure 5f**). As shown in **Figure 5e and f**, the most significant gene set (Set 1) linked to the lLG prediction exhibited conspicuous enrichment in a consistent set of terms associated with neuroactive signaling. These terms encompassed critical aspects such as neuroactive ligand-receptor interaction, chemical synaptic transmission, calcium ion binding, and behaviors elicited by internal or external stimuli. These processes can influence the configuration of neural circuits, their information processing mechanisms, and neural plasticity within the hippocampus^69^, which could support and reinforce the functional gradient along the long axis. Furthermore, gene expression pertaining to the regulation of anatomical structure morphogenesis and neural development plays an important role in predicting variance in the lLG. This observation mirrored the prior discovery stemming from the gene set associated with predicting anatomical positions along the human hippocampal anterior-posterior axis (y coordinates)^5^. Importantly, molecular processes implicated in the regulation of stress hormones, i.e., positive regulation of cortisol secretion, also displayed variations along the lLG axis. This finding aligned with established literature illustrating the hippocampus’s critical involvement in regulating stress-related cortisol activity^70,71^ and its high density of glucocorticoid receptors^72,73^. Consequently, our findings offer a potential link between the regulation/targeting of stress-related cortisol activity and the hippocampal linear long-axis connectome gradient. This novel perspective holds significance in exploring the impact of adversity or stress responses on neuroplasticity^74,75^ and neurodevelopment^76,77^, as well as their implications for relevant brain disorders such as depression^78,79^. Detailed information on gene annotation and enrichment can be found in Supplementary Data 1. Notably, analogous outcomes for enrichment analysis of biological pathways were also derived from the gene list within Set 2. This analysis revealed similar clusters of terms associated with neuroactive signaling, neural development, and hormone activity (see **Extended Data Fig. 10a and b**).

To ascertain whether the noteworthy gene set related to lLG exhibits enrichment specifically within the hippocampus and during the developmental period under investigation, genes identified through the above analysis were compared against developmental expression profiles from the BrainSpan dataset (http://www.brainspan.org/) by employing the developmental specific expression analysis (SEA) tool^67^. The results underscored that the important gene expression (Set 1) exhibited enrichment within the hippocampus during mid/late childhood, adolescence, and young adulthood. This enrichment was notably accentuated in the mid/late childhood period (depicted in the left and middle panels of **Figure 5g**), which aligned with the fastest developmental period in the hippocampal lLG (see left panel in **Figure 2b**). Comparable results were also obtained from the developmental enrichment analysis of the Set 2 gene list (see **Extended Data Fig. 10c**). These outcomes potentially imply that the previously identified molecular processes involving neuroactive signaling, stress hormone regulation, and neural growth likely contribute to the development of the hippocampal linear long-axis gradient in youth. Moreover, the genes within Set 1 were also found to be enriched within the hippocampus during the period spanning from late/mid fetus to early infancy. Considering existing evidence that the long-axis functional gradient might not be fully formed at birth^80^, these findings may suggest that the differentiation of the hippocampal long-axis could experience significant development during this particular period. Furthermore, these genes were found to be expressed in the cortex during early infancy, childhood, adolescence, and young adulthood (see the right panel of **Figure 5g**). This observation suggests that the molecular processes related to these genes could also play a significant role in the maturation of the cortex during sensitive developmental periods.

These micro-level results highlight the potential role of neuroactive signaling, stress hormone regulation, and neural growth in enhancing the development of the hippocampal linear long-axis gradient in youth. This observation suggests that experiences and environmental factors including adversity or stress influencing hippocampal neural activity and neural plasticity might drive the substantial maturation of the dominant functional organizational principle within the hippocampus during this sensitive period of development. These findings potentially offer insights for designing neurobiologically informed interventions that can be effectively delivered to improve hippocampal developmental outcomes or treat relevant developmental disorders.

## Discussion

Like many natural systems, both the hippocampus and the cortex, depend on a set of fundamental organization principles to process information, regulate behavior, and facilitate various intricate functions. We integrated cutting-edge methods from neuroanatomy, transcriptomics, physics, machine learning, and cognitive neuroscience to unveil the intricate organization of the hippocampus-cortex interactions in human and its developmental significance. Here, we provide compelling evidence for the presence of a dual long-axis on gradients within the human hippocampus by using two independent developing cohort datasets. These gradients were evident in terms of hippocampal-cortical connectivity, hippocampal geometric eigenmodes, and their roles in supporting cortical hierarchies, indicating that the organizational principles are not only evident in how the hippocampus communicates with the neocortex but also in its structural properties and influence on the hierarchical organization of high-order functions. Our results challenge classical views on hippocampal long-axis differentiation, which have predominantly (except^81^) identified a consistent monotonic gradient across multiple organizational traits^1,2,4–6,24^. We propose that the organization and function of the hippocampus are more complex and nuanced than previously thought. This has potential implications for our comprehension of memory, spatial navigation, and other associated processes.

Notably, we found that dual long-axis gradients, particularly the quadratic long-axis gradient within the hippocampus, play a critical role in its integration into cortical hierarchical processing. This implies that the structure and functioning of the hippocampus are closely linked to how the cortex processes and organizes information. Given the greater dominance of hierarchical processing in humans^9^ and macaques^10^ in contrast to other species such as marmosets^11^ or rodents^32^, these intriguing findings call into question the wide belief of the hippocampus as an evolutionarily conserved structure concerning its structural and functional organization^1,13,14^. These results may begin a novel paradigm for exploring these organization axes of the hippocampus, now incorporating the quadratic long-axis gradient. This paradigm underscores the need for meticulous cross-species comparison studies on the hippocampus’s integration into cortical systems to gain deeper insights into its role in cognition and brain function and its potential implications for various brain disorders.

The geometric constraints on the dual long-axis gradients continue to diminish, potentially contributing to the maturation of cognitive function such as executive function as we demonstrated. Molecular processes underlying the maturation of the hippocampal linear long-axis gradient may be associated with neuroactive signaling, stress hormone regulation, and neural growth. Together these findings enrich our comprehension of the development of hippocampal organizational principles on functional, structural, and molecular dimensions in youth and their importance in cognitive development. They provide insights into the reorganization of hippocampal integration with large-scale cortical hierarchy systems and signify significant developmental milestones during the transition to adolescence and adulthood, reflecting the dynamic reorganization of neural circuits and cognitive processes associated with these two inflection points.

The restructuring of connectivity between both the frontoparietal and ventral attention systems as a signature of the dual long-axis reorganization plays a critical role in the hippocampus’s contribution to the maturation of the cortical hierarchy. This phenomenon may be attributed to the potentially significant role of these two systems in driving the reorganization of the cortical functional connectome in youth, and facilitating across-system communication that potentially coordinates both bottom-up and top-down information flow, as evidenced recently^60,62,82,83^. Our findings on the reorganization of connectivity between these cortical systems and the long-axis of the hippocampus indicated how the ongoing maturation of the hippocampus, with support from its critical organizational axes, integrates with continuously maturing large-scale cortical systems. These integration changes promote the maturation of the cortical hierarchy and potentially enhance efficient communication between cortical systems, motivating future research to elucidate potential cognitive and behavioral implications.

The geometric eigenmodes derived from both neocortical and non-neocortical structures have recently been demonstrated to be a more concise and precise representation of their macroscale functional activity compared to the connectome-based method^8^. Furthermore, multiple recent studies have demonstrated that wave dynamics constrained by geometry and distance-dependent connectivity may have a dominant influence on spatiotemporal brain activity^8,62,84^. Therefore, the reduction trajectories in structure-function coupling between hippocampal geometry and dual long-axis functional gradient suggest the development of long-range connectivity and that the dominance of wave dynamics within the hippocampus may decrease during its maturation process. We inferred that this decoupling process may enable efficient communication both within and outside the hippocampus, bolstering the complexity of hierarchical cortical processing. Ultimately, this process may play a role in advancing high-order cognitive functions. Our findings revealed a link between the structure-function long-axis reorganization and the core EF performance, providing further support for this inference. These relationships varied across the dual long-axis gradients, suggesting distinct functional implications for the linear long-axis gradient and the quadratic long-axis gradient. The functional and developmental significance of the wave-like propagation of hippocampal functional activity along the dual long-axis gradient, based on techniques such as optical flow^85^ or quasiperiodic pattern identification^84^, are warranted in future.

A consistent spatial axis of neurodevelopmental variability in both myelin content and functional gradient profiles exists during this sensitive period. This spatial axis closely aligns with spatial variations in myelin content itself, a widely recognized factor known to restrict neural plasticity^55,56,86,87^. The existence of critical periods with increased plasticity for hippocampus-dependent learning and memory has been discovered and has gained strong support in recent years^35–38^. Additionally, the formation of myelin and myelin-related neurite outgrowth inhibitor (Nogo) receptor signaling has been identified as a constraining or closure mechanism for the critical period in the primary sensory cortex^55,56^, and Nogo receptors were also found to restrict synapse formation in the developing hippocampus^88^. Our findings, which indicated that myelin regulates the developmental plasticity of hippocampal functional gradients, align with these discoveries, and potentially imply that myelin also plays a role in regulating the critical period plasticity of hippocampal development in youth. Our discoveries contribute to the understanding of how plasticity is distributed across hippocampal regions at various developmental stages. This knowledge is critical for further comprehending the effects of diverse environmental, experiential, and neurobiological factors that converge on neurodevelopmental outcomes during periods of heightened plasticity. Our study has the potential to guide future research and therapeutic strategies aimed at enhancing cognitive development and addressing neurological conditions. Future studies could consider combining pharmacological or chemogenetic approaches with neuroimaging techniques to establish connections between cellular-or molecular-level plasticity-regulating mechanisms and noninvasive human neuroimaging^89–91^. This integrated approach can provide further insights into specific neurobiological mechanisms underlying critical periods in the development of the human hippocampus.

Of note, we demonstrated the potential role of neuroactive signaling, stress hormone regulation, and neural growth in enhancing the development of the hippocampal linear long-axis gradient in youth. Specifically, neuroactive ligand-receptor interactions and neural growth are two key molecular processes involved, encompassing neurotransmitter and growth factor binding to receptors, which regulate neural activity and growth. The varied expression of these processes along the hippocampal long axis may lead to variations in neural circuitry and connectivity. Notably, we found that the regulation of stress hormones, i.e., positive regulation of cortisol secretion, also displayed variations along the linear long-axis gradient. This novel perspective emphasizes a potential association between the regulation/targeting of stress-related cortisol activity and the long-axis differentiation of the hippocampal connectome. It offers a new mechanistic framework for comprehending the well-established role of the hippocampus in regulating stress-related cortisol activity^70,71^ and how it is influenced by this activity, given the high density of glucocorticoid receptors within the hippocampus^72,73^. This framework can be substantiated through the design of more rigorous experiments based on animal models^92^ in future research. Future studies can also investigate how mental training programs for humans, known to benefit stress reduction^93,94^ influence the linear long-axis gradient of the hippocampus. Adversity and stress responses are widely recognized risk factors for impaired neuroplasticity^74,75,95^ and neurodevelopment^76,77,96^. These factors also have significant implications for relevant brain disorders such as depression^78,79^. Future research can investigate how stress-induced clinical disorders or experiences of youth adversity disrupt or influence the hippocampal linear long-axis gradient. On the other hand, we did not obtain a robust gene list to account for the hippocampal qLG based on our linear model, and we inferred that this organization principle of the hippocampus, which is not as evolutionarily conserved as the lLG, may be regulated by gene expression in a more complex and nonlinear manner.

We noted that the developmental analysis was conducted based on cross-sectional data (latest release of the HCP-D). Future studies based on longitudinal datasets could further characterize the development of dual long-axis gradients within individuals. Though we leveraged the latest release of the HCP-D dataset, it did not encompass measures of pubertal hormones.

Consequently, the pubertal stage we referred to in this paper was broadly defined. This constraint hindered our capacity to conclusively determine whether hippocampal dual long-axis gradients were affected by puberty or hormone levels. Notably, rodent studies have shown that pubertal hormones can affect neurobiological mechanisms of plasticity in the hippocampus^97^, such as inhibitory signaling^98^. This underscores the need for future research in this area. The present study demonstrated that in youth, the dual long-axis differentiation of the functional connectome and geometric eigenmodes forms and undergoes significant development. Future studies can be conducted to study the formation and developmental trajectories of dual long-axis organization and the cognitive function and behavior involved during infancy and early childhood. Indeed, the well-established cytoarchitectonic variation within the hippocampus has long been recognized as a critical organizational principle. This often leads to the division of the hippocampus into separate subfields, such as the dentate gyrus (DG), cornu ammonis (CA), and subiculum (Sub). Investigating the distinct functional gradient characteristics of these subfields and their maturation process is a promising avenue for future research.

## Data availability

All data are available in the main text or the supplementary materials. Segmentations and mid-thickness surfaces of the hippocampus, the sampled BOLD time series on the generated mid-thickness surfaces and cortical parcels, for both the HCP-D and CBD datasets, are available upon request. The original and preprocessed HCP-D data, after meeting eligibility requirements, can be accessed here: https://humanconnectome.org/study/hcp-lifespan-development.

## Code availability

The code used will be available on GitHub upon this paper is accepted.

## Acknowledgments

Shuyu Li is supported by NSFC (32271146) and the Startup Funds for Top-notch Talents at Beijing Normal University. Yong He is supported by NSFC (82021004). Qiongling Li is supported by NSFC (82202245). Qi Dong is supported by NSFC (31521063). Sha Tao, Yanpei Wang, Daoyang Wang, Mingming Hu, and Zhiying Pan are supported by NSFC (31521063) & Beijing Municipal Science & Technology Commission (Z15110000391512). Dr. Zuo is supported by the STI 2030 - the major projects of the Brain Science and Brain-Inspired Intelligence Technology (2021ZD0200500), the Start-up Funds for Leading Talents at Beijing Normal University.

## Author contributions

Shuyu L., X.N. Z., Yong H., and D. L. conceptualized the work. D. Z. contributed to hippocampal segmentation and mid-thickness surface generation. Yirong H. and X. D. conducted the quality control of segmentations and surfaces of the hippocampus for HCP-D data. Shaoxian L. and S. B. conducted this quality control for CBD data. D. Z. and Q. L. performed MR image preprocessing for CBD data. D. Z. and Q. L. conducted structural and functional feature mapping and data analysis. D. Z., X.N. Z., Shuyu L., and Q. L. conducted data interpretation. D. Z. and Q. L. performed data visualization. W. M., Y. W., Shuping. T., J. G., S. Q., Sha T., Q. D. and Yong H. contributed to MR images acquisition of CBD data. Sha T., D. W., Y. W., M. H., and Z. P. ran the cohort and conducted all arrangements of CBD data. Y. X. and L. S. organized these data. X. L., T. Z., X. C. and T. L. performed the quality control of MR images for CBD data. D. Z., X.N. Z., Yong H., Shuyu L., Q. L. and Yirong H. wrote the paper.

## Competing interests

The authors declare no competing interests.

---

**Extended Data Fig. 1:**
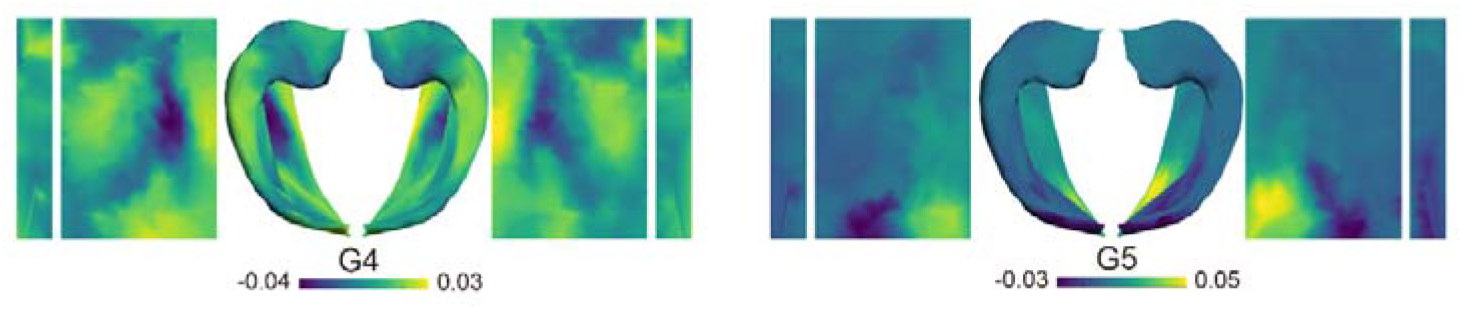
Topographic pattern of the 4^th^- and 5^th^-order hippocampal-cortical connectivity gradient.

**Extended Data Fig. 2:**
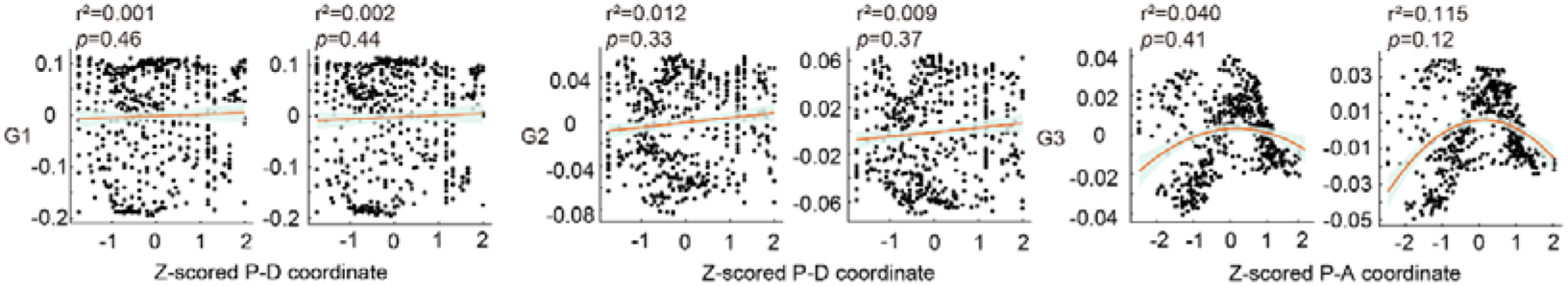
Relationships between each of the first three hippocampal gradients and the alternative geodesic axis.

**Extended Data Fig. 3:**
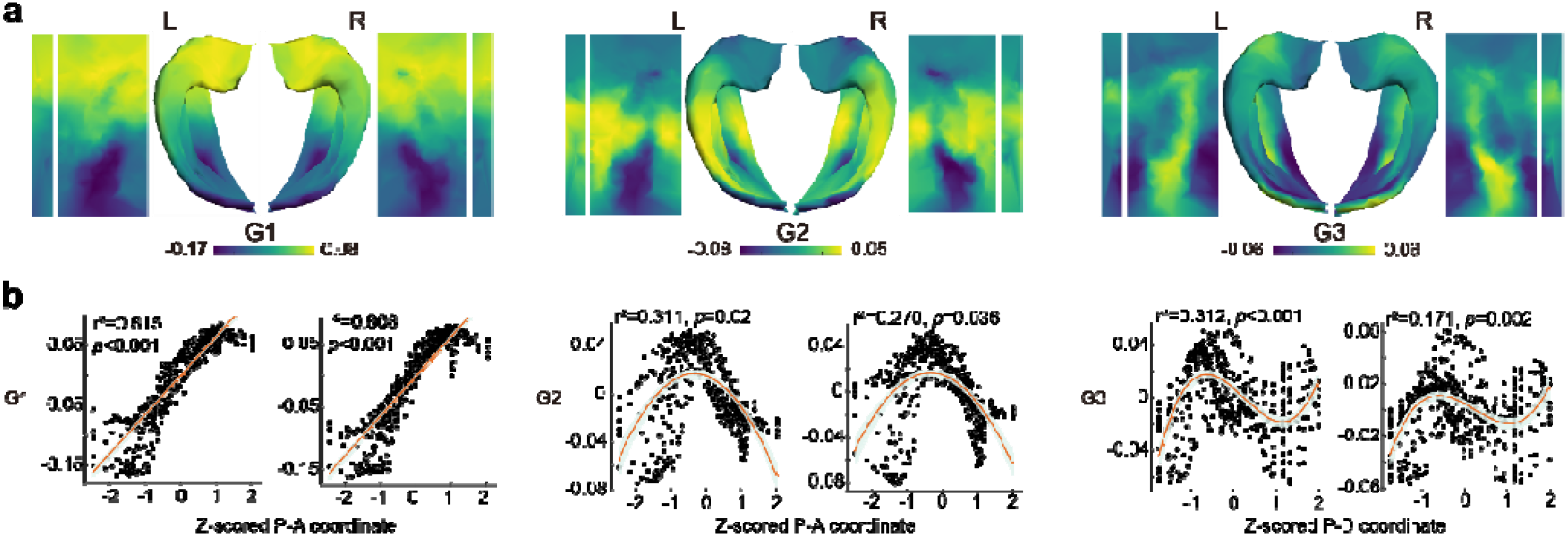
Replicated results for the first three hippocampal-cortical functional gradients in the CBD dataset.

**Extended Data Fig. 4:**
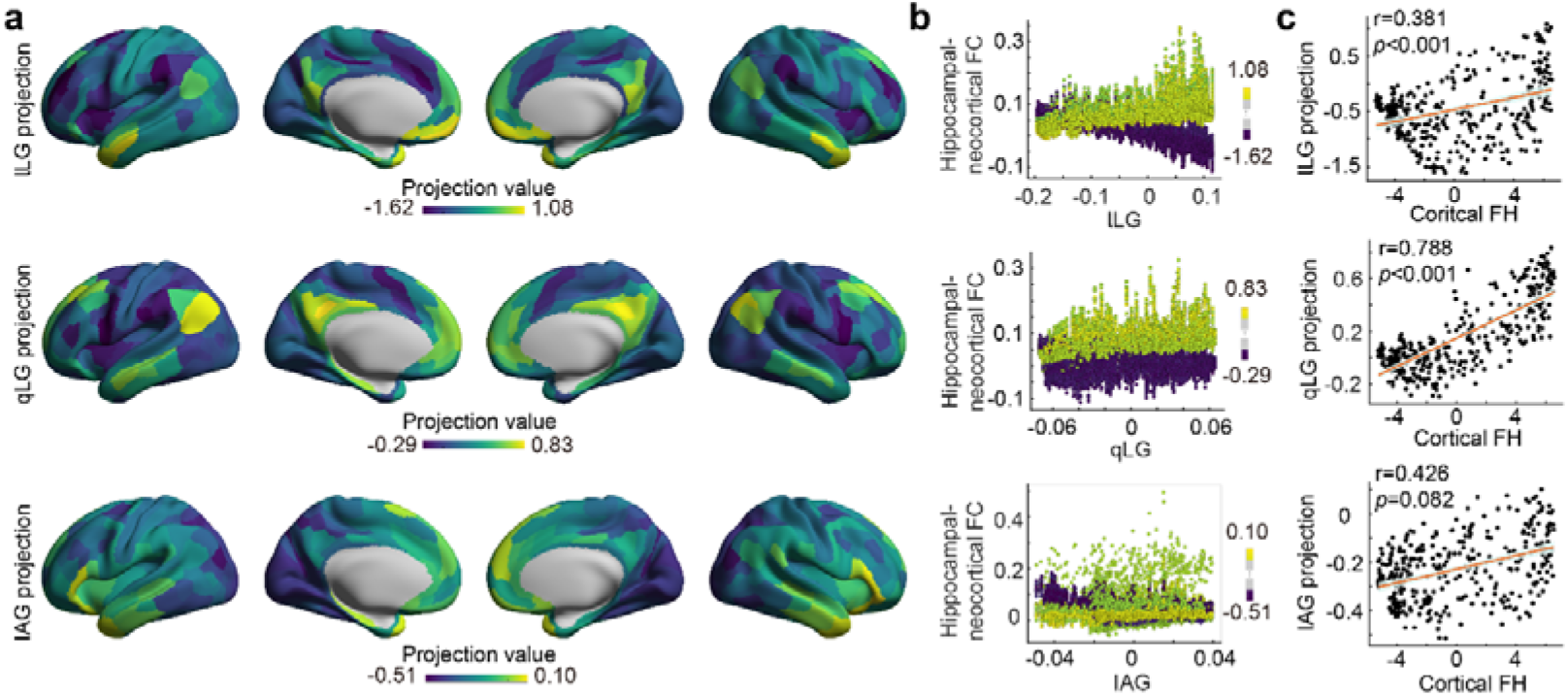
Cortical projection of the right hippocampal gradients in youth. **a**, Group-averaged cortical projection of the right hippocampal gradients. **b**, Relationship between the right hippocampal gradients and the neocortical-hippocampal FCs corresponding to the cortical regions that exhibit the top 5% and bottom 5% projection values. Each data point within the diagram signifies an individual vertex located on the hippocampal surface. The color indicates the projection values. **c**, Correlation between the projection pattern of specific gradients in the right hippocampus and the cortical FH.

**Extended Data Fig. 5:**
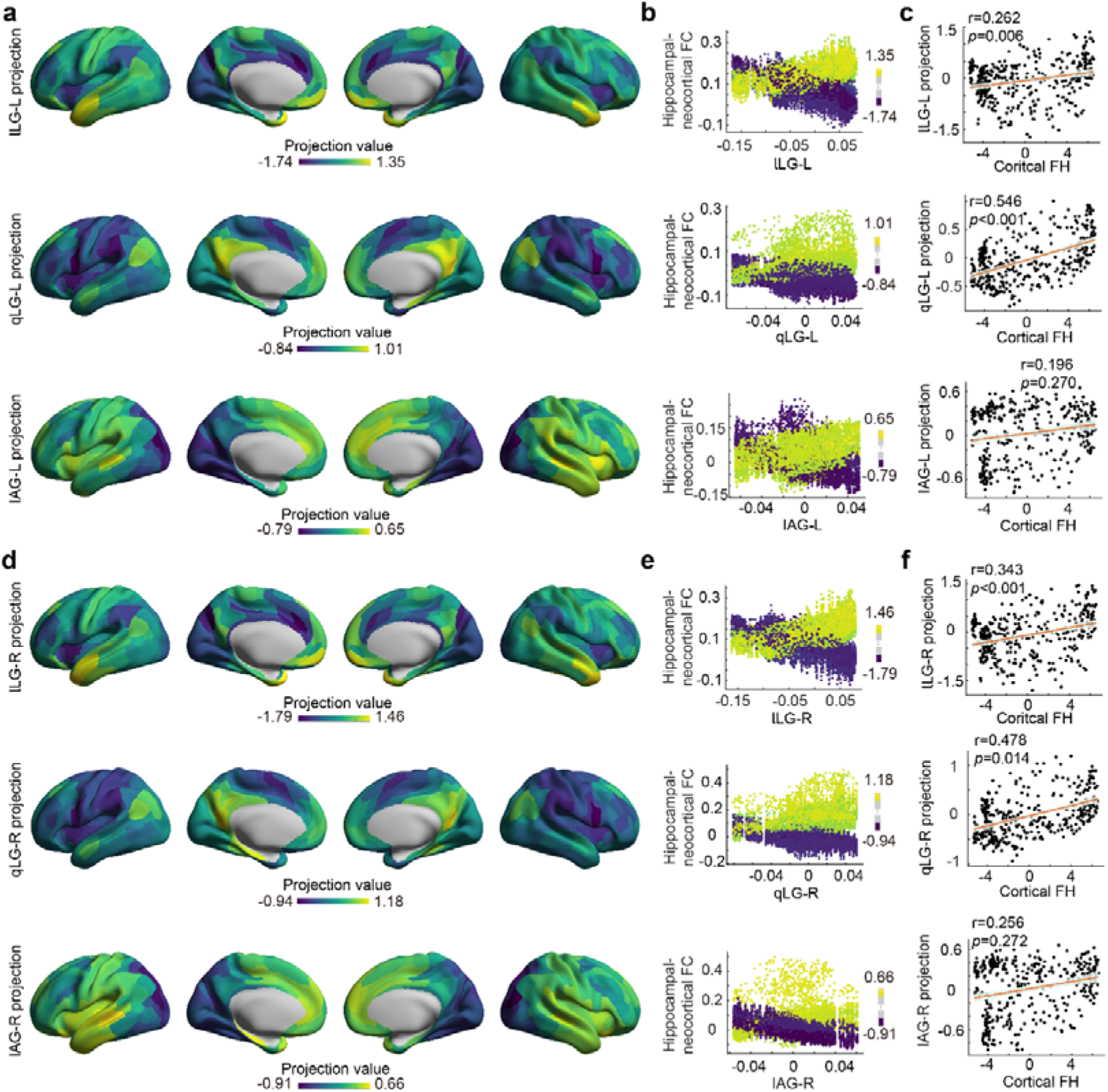
Validation results for cortical projections of the first three hippocampal gradients in the CBD dataset. **a, b,** and **c** correspond to the results of the left hippocampus while **d, e,** and **f** correspond to those of the right hippocampus.

**Extended Data Fig. 6:**
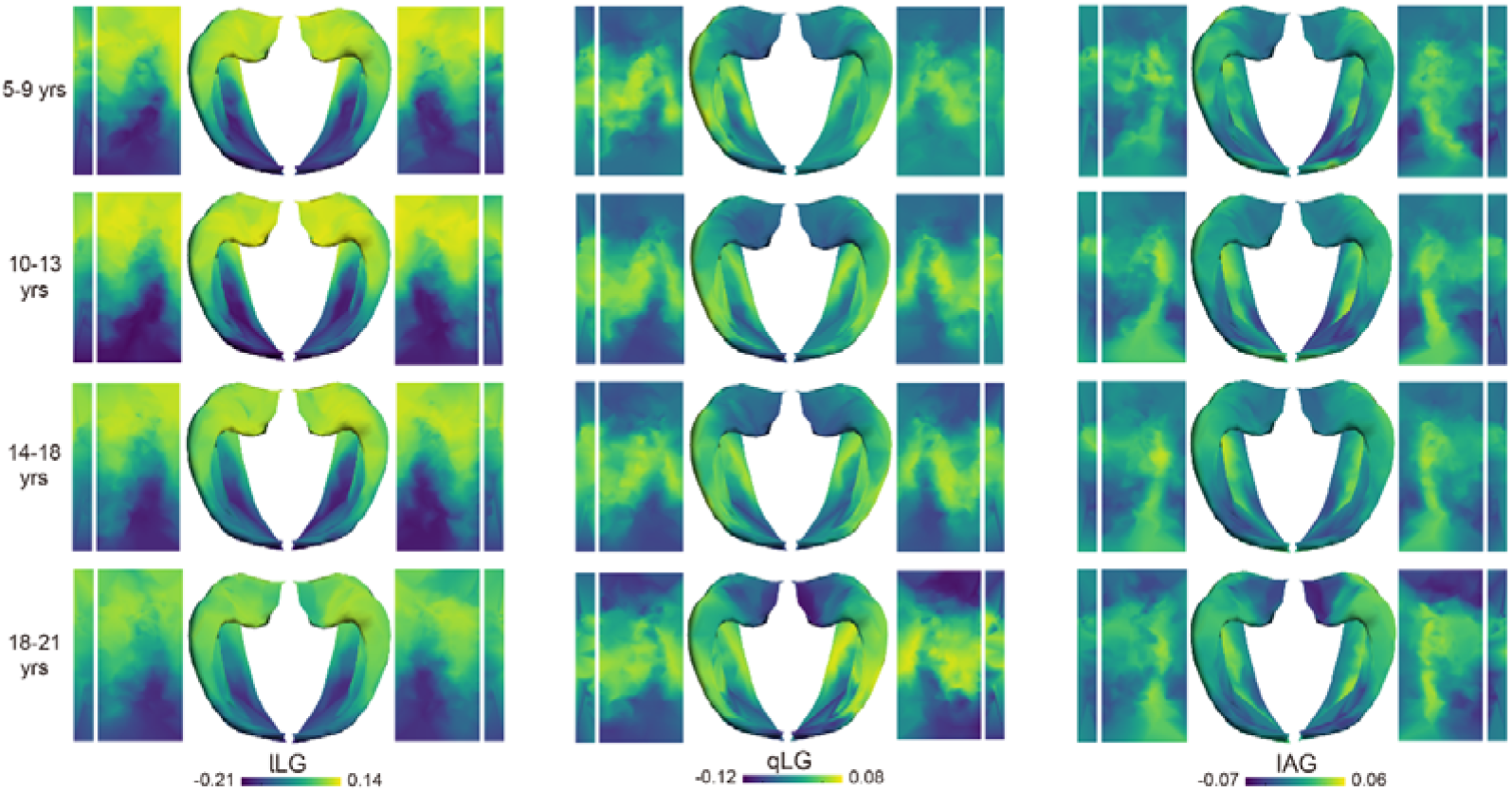
Topographic pattern of the first three gradients presented within four age-specific groups (5-9, 10-13, 14-17, and 18-21 years).

**Extended Data Fig. 7:**
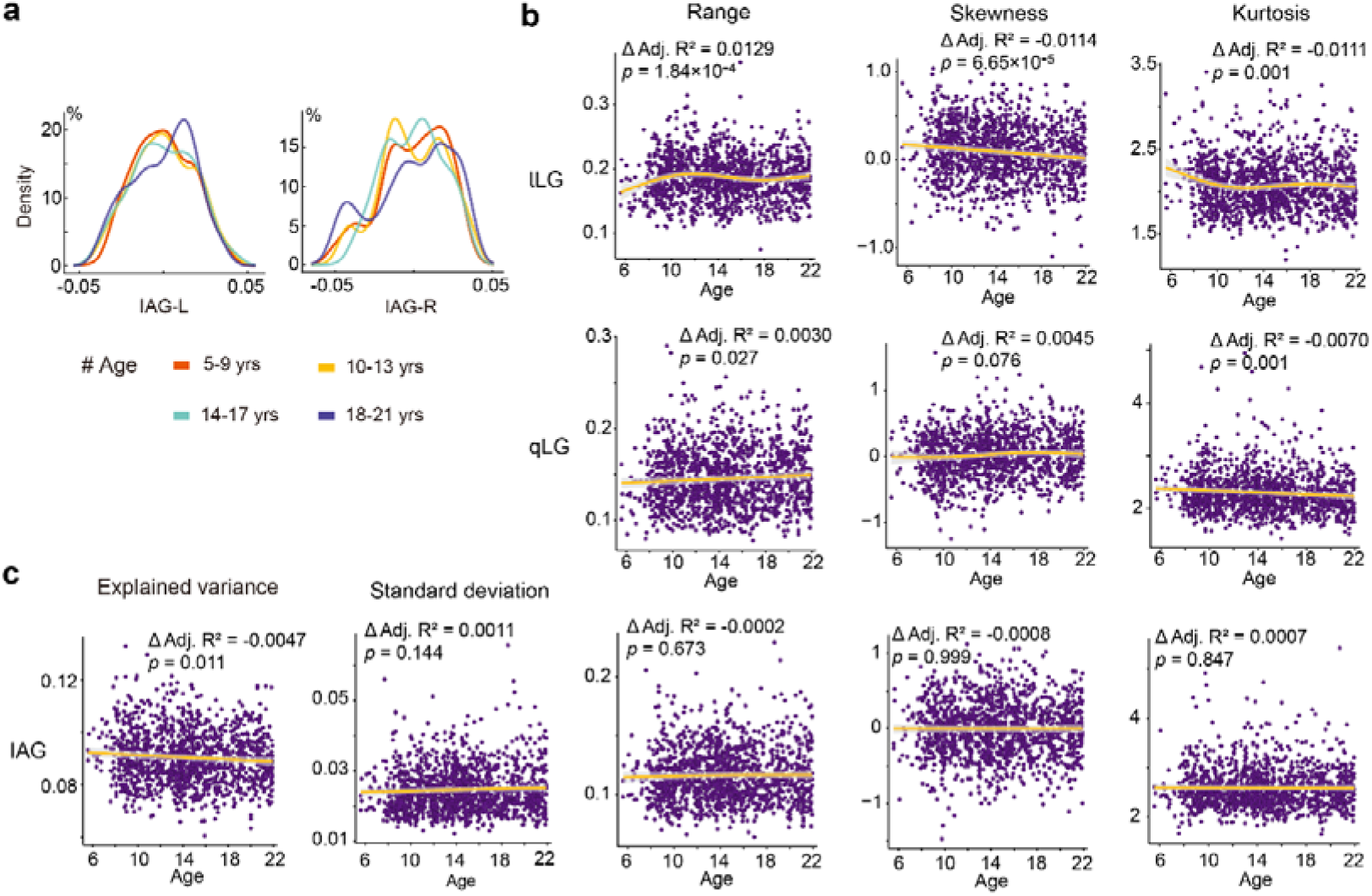
Additional results regarding the development of the distribution characteristics of the first three gradients. **a,** Density estimation of the iso-allocortical gradient (IAG) within four age-specific groups (5-9, 10-13, 14-17, and 18-21 years). **b**, Developmental progression of the range, skewness, and kurtosis of the dual long-axis gradients. **c**, Developmental progression of the explained variance, standard deviation, range, skewness, and kurtosis of the IAG.

**Extended Data Fig. 8:**
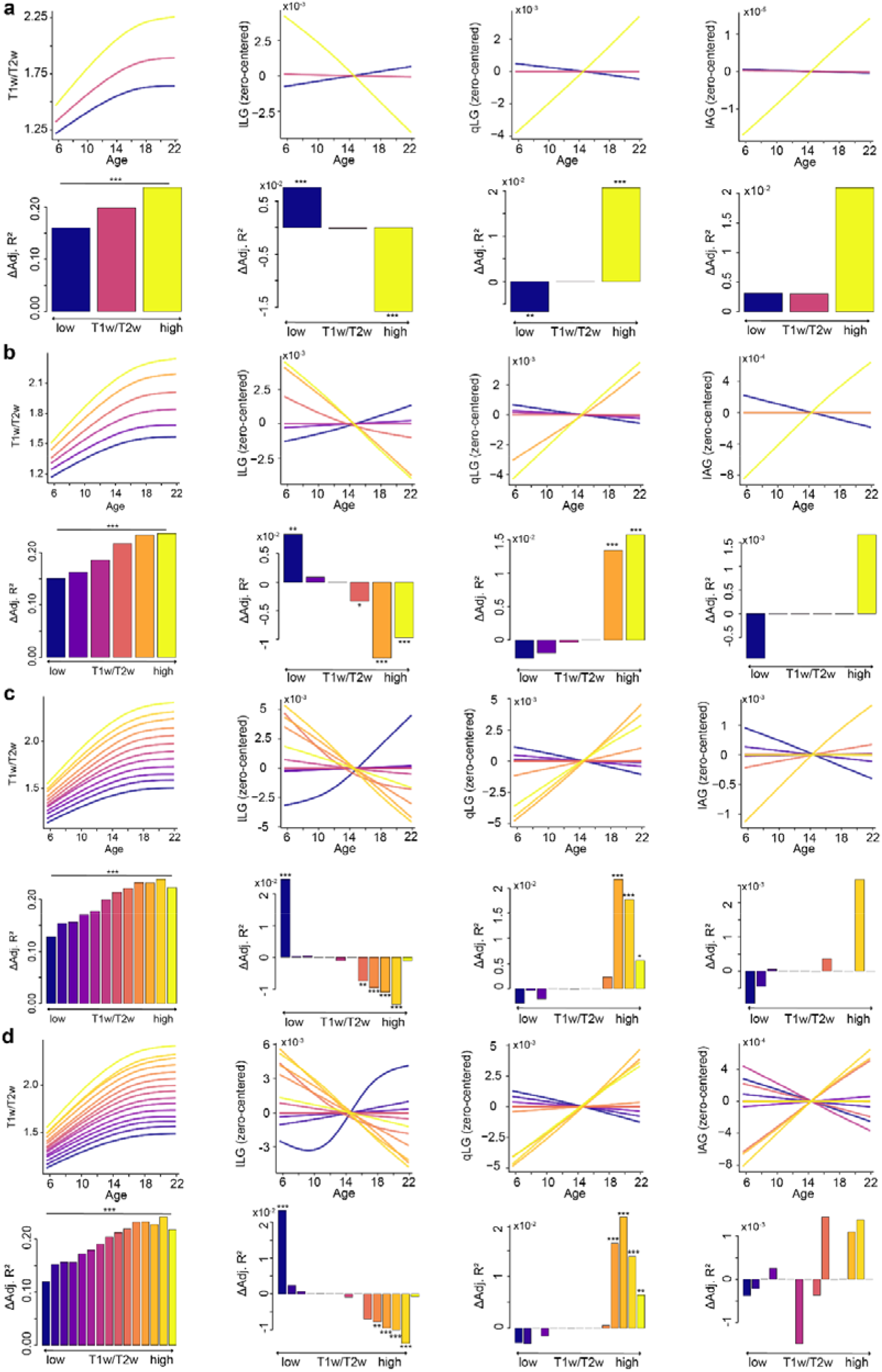
Spatial and temporal correspondence between the refinement of myelin content and the maturation of hippocampal functional gradients is replicated with various bins (3, 6,12 and 15) used for partitioning the hippocampal mid-thickness surface.

**Extended Data Fig. 9:**
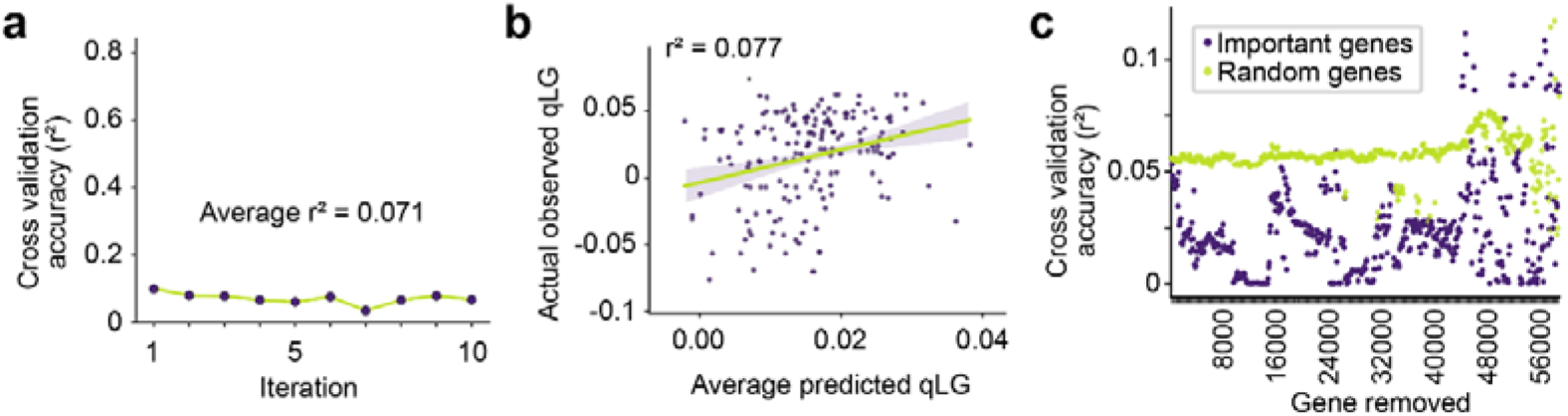
Prediction of the hippocampal qLG using ten separate 10-fold cross-validated LASSO-PCR models based on transcriptomic data, with a feature deconstructing process to explain gene importance.

**Extended Data Fig. 10:**
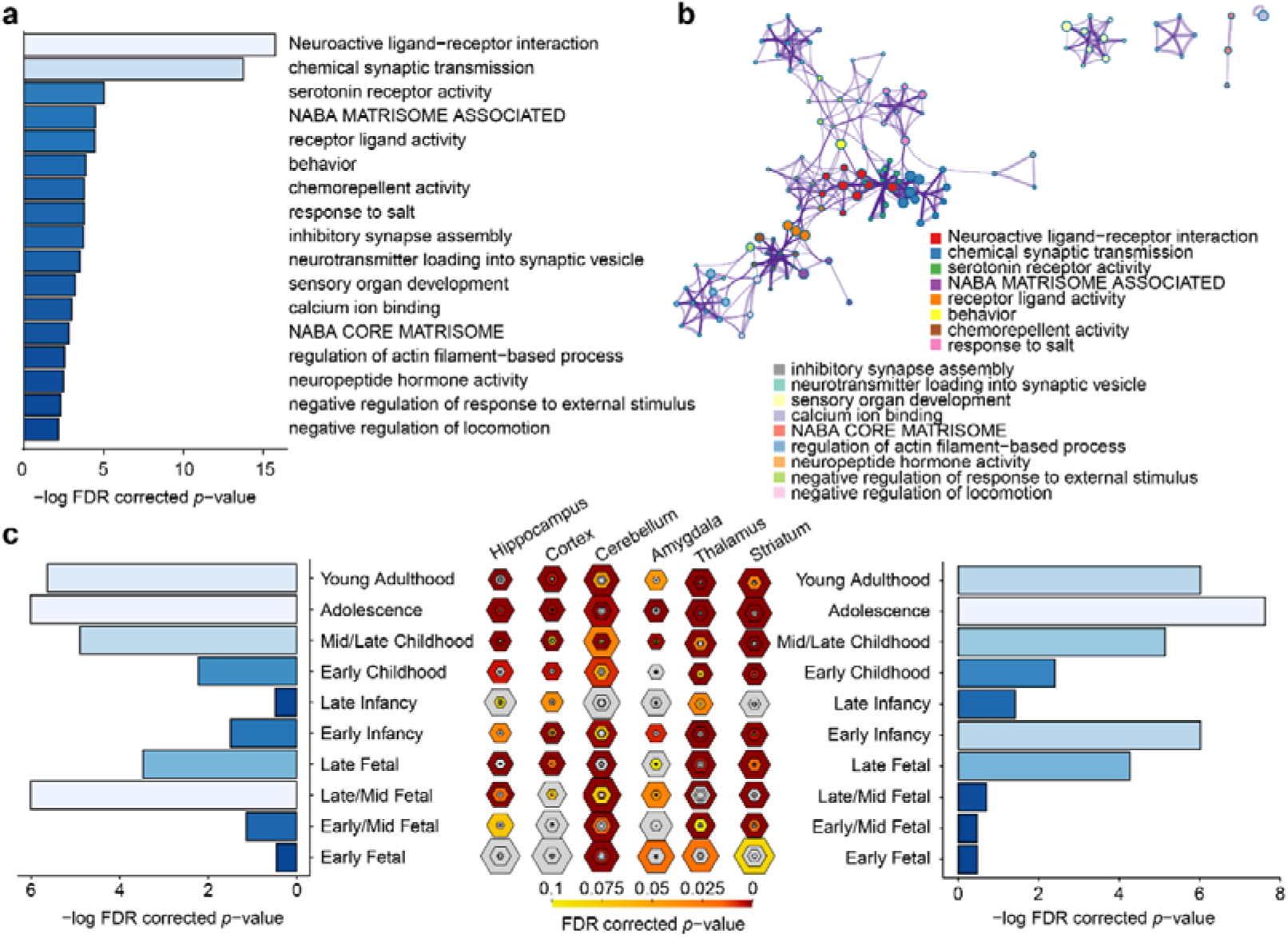
Biological pathways and development enrichment analysis on the important gene list of Set 2 for the lLG.

**Extended Data Table 1.** Δ Adjusted R^2^ and FDR-corrected *p* value for nine bins along the group-averaged T1w/T2w axis.

| Bin | T1w/T2w |  | G1 |  | G2 |  | G3 |  |
| --- | --- | --- | --- | --- | --- | --- | --- | --- |
| | $\Delta$ Adj. $R^2$ | $p$ value | $\Delta$ Adj. $R^2$ | $p$ value | $\Delta$ Adj. $R^2$ | $p$ value | $\Delta$ Adj. $R^2$ | $p$ value |
| 1 | 0.1409 | <1E-16 | 0.0131 | 0.0001 | -0.0046 | 0.0169 | -0.0008 | 0.6378 |
| 2 | 0.1549 | <1E-16 | 0.0015 | 0.1059 | -0.0026 | 0.0650 | 2.88E-06 | 0.8819 |
| 3 | 0.1662 | <1E-16 | 0.0002 | 0.3013 | 3.0E-7 | 0.8673 | -5.38E-07 | 0.8819 |
| 4 | 0.1800 | <1E-16 | -8.1E-7 | 0.5489 | 7.8E-8 | 0.5518 | 1.35E-06 | 0.8819 |
| 5 | 0.2092 | <1E-16 | -0.0017 | 0.1059 | -7.2E-6 | 0.5493 | 2.33E-06 | 0.8819 |
| 6 | 0.2197 | <1E-16 | -0.0070 | 0.0077 | -1.1E-6 | 0.7359 | 0.0001 | 0.8508 |
| 7 | 0.2313 | <1E-16 | -0.0102 | 0.0005 | 0.0076 | 0.0030 | -1.93E-06 | 0.9729 |
| 8 | 0.2390 | <1E-16 | -0.0174 | 1.6E-5 | 0.0228 | <1E-16 | 0.0009 | 0.6378 |
| 9 | 0.2298 | <1E-16 | -0.0037 | 0.0154 | 0.0106 | 0.0005 | -2.39E-07 | 0.8819 |

## Notes

### Competing Interest Statement

The authors have declared no competing interest.

